# Taxonomic distribution of metabolic functions underpins nutrient cycling in *Trichodesmium* consortia

**DOI:** 10.1101/2023.03.15.532517

**Authors:** Coco Koedooder, Futing Zhang, Siyuan Wang, Subhajit Basu, Sheean T. Haley, Nikola Tolic, Carrie D. Nicora, Tijana Glavina del Rio, Sonya T. Dyhrman, Martha Gledhill, Rene M. Boiteau, Maxim Rubin-Blum, Yeala Shaked

**Affiliations:** The Fredy and Nadine Herrmann Institute of Earth Sciences, Hebrew University of Jerusalem, Jerusalem, Israel; The Interuniversity Institute for Marine Sciences in Eilat, Eilat, Israel; Israel Oceanographic and Limnological Research, Haifa, Israel; Microsensor Research Group, Max Planck Institute for Marine Microbiology, Bremen, Germany; Lamont-Doherty Earth Observatory, Columbia University, New York, NY, United States; Biological Sciences Laboratory Pacific Northwest National Laboratory, Richland, WA, United States; Joint Genome Institute, Lawrence Berkeley National Laboratory, Berkeley, CA, United States; Department of Earth and Environmental Sciences, Columbia University, New York, NY, United States; GEOMAR, Helmholtz Center for Ocean Research, Kiel, Germany; Environmental Molecular Sciences Laboratory Pacific Northwest National Laboratory, Richland, WA, United States; College of Earth, Ocean, and Atmospheric Sciences, Oregon State University, Corvallis, OR, United States

**Keywords:** *Trichodesmium*, metagenomes, associated bacteria, siderophore biosynthesis, vitamin B, denitrification, dissimilatory nitrate fixation to ammonia, phosphonate, nitrogen fixation

## Abstract

The photosynthetic and diazotrophic cyanobacterium *Trichodesmium* is a key contributor to marine biogeochemical cycles in the subtropical-oligotrophic oceans. *Trichodesmium* forms colonies that harbor a distinct microbial community, which expands their functional potential and is predicted to influence the cycling of carbon, nitrogen, phosphorus and iron (C, N, P, and Fe). To link key traits to taxa and elucidate how community structure influences nutrient cycling, we assessed Red Sea *Trichodesmium* colonies using metagenomics and metaproteomics. This diverse consortium comprises bacteria that typically associate with algae and particles, such as the ubiquitous *Alteromonas macleodii,* but also lineages specific to *Trichodesmium*, such as members from the order Balneolales. These bacteria carry functional traits that would influence resource cycling in the consortium, including siderophore biosynthesis, reduced phosphorus metabolism, vitamins, denitrification, and dissimilatory-nitrate-reduction-to-ammonium (DNRA) pathways. Denitrification and DNRA appeared to be modular as bacteria collectively completed the steps for these pathways. The vast majority of associated bacteria were auxotrophic for vitamins, indicating the interdependency of consortium members. *Trichodesmium* in turn may rely on associated bacteria to meet its high Fe demand as several lineages can synthesize the photolabile siderophores vibrioferrin, rhizoferrin, and petrobactin, enhancing the bioavailability of particulate-Fe to the entire consortium. Our results highlight that *Trichodesmium* is a hotspot for C, N, P, Fe, and vitamin exchange. The functional redundancy of nutrient cycling in the consortium likely underpins its resilience within an ever-changing global environment.

**Importance:** Colonies of the cyanobacteria *Trichodesmium* act as a biological hotspot for the usage and recycling of key resources such as C, N, P and Fe within an otherwise oligotrophic environment. While *Trichodesmium* colonies are known to interact with a unique community of algae and particle-associated microbes, our understanding of the taxa that populate these colonies and the gene functions they encode is still limited. Characterizing the taxa and adaptive strategies that influence consortium physiology and its concomitant biogeochemistry is critical in a future ocean predicted to have increasing particulate fluxes and resource-depleted regions.

## Introduction

*Trichodesmium* spp. are a globally relevant group of cyanobacteria that can form surface blooms visible from space (1). Owing to their high abundance and capability to fix both carbon and nitrogen, they are considered key players in the biogeochemical cycling of C and N within the oligotrophic tropical and subtropical oceans (2–5). A key trait underpinning the functionality of *Trichodesmium* is that its filaments can cluster together to form large colonies of 1-2 mm in size. The formation of colonies allows *Trichodesmium* to capture dust that is deposited on the ocean surface and *Trichodesmium* is reported to actively select for Fe and P-rich particles and shuffle them to their colony core (6–8). Colony formation hereby adds an intriguing spatial component where *Trichodesmium* spp. is predicted to act as a biological hotspot not only for C and N, but also for the uptake and recycling of key limiting nutrients such as Fe and P.

*Trichodesmium* colonies harbor a unique microbial consortium of epibiotic bacteria. Amplicon sequencing has shown these associated bacteria are distinct from populations in the surrounding oligotrophic surface waters (9–11). At a genetic level, the associated microbial community was found to enrich the total functional potential of the collective colony beyond the traits of N_2_ and C fixation (12–14). By enriching the genetic repertoire of metabolic functions, or traits, related to the nutrients C, N, Fe, and P, consortia members were predicted to influence the internal cycling of these nutrients (14–17). To understand the biogeochemical contributions of *Trichodesmium* within the global ocean it is therefore critical to look beyond the metabolic functions present in *Trichodesmium* and towards the processes taking place within the larger microbial community. While metagenomic studies of *Trichodesmium* colonies in the Atlantic (13) and the Pacific Oceans (12, 17, 18) highlighted key functional traits within the consortium, only one study linked pathways to the associated bacteria from 8 metagenome-assembled bins (13). Expanding these studies, and characterizing how functional traits are distributed across different taxa is critical to understanding the role of *Trichodesmium* associated bacteria in consortium physiology and its concomitant influence over the cycling of key resources such as C and N.

Recent evidence from the Red Sea highlights the unique ability of *Trichodesmium* consortia members to acquire particulate Fe from dust through the production of Fe-chelating siderophore molecules (7, 8, 16, 19, 20). Siderophores dissolve particulate Fe from dust and are predicted to increase its bioavailability for both *Trichodesmium* and its associated bacteria (16, 21, 22). In this manner, *Trichodesmium* spp. interacts with their associated bacteria in order to meet the high Fe-requirements for the processes of photosynthesis and N_2_ fixation (23). This finding is particularly relevant in light of climate model simulations, which predict an increase in regional dust emissions in areas such as the Red Sea where *Trichodesmium* is known to occur (24, 25). While *Trichodesmium* blooms are a re-occurring phenomenon in this area (26), it represents an under sampled region where the taxonomic composition of its consortium has not been explored genetically. Furthermore, the presence and taxonomic distribution of siderophore biosynthesis pathways within *Trichodesmium* colonies has not yet been interrogated in depth. Understanding which consortium members positively enhance the bioavailability of Fe within the *Trichodesmium* consortium will help elucidate key interactions and community dynamics taking place within the colony that influence the ability of *Trichodesmium* to fix C and N. This information is critical to evaluating *Trichodesmium* contributions to C and N cycling in the future ocean.

In this study, we conducted an in-depth metagenomic and proteomic study of *Trichodesmium* colonies collected from the Red Sea. To elucidate how the community structure influences the functional dynamics of the *Trichodesmium* consortium we identified the different taxa making up the *Trichodesmium* consortium and examined the presence of functional traits involved in nutrient cycling. We probed our dataset for genes, proteins and pathways involved in the uptake and metabolism of N, P, Fe, and vitamins. Finally, we contextualize these findings by comparing them to previous metagenomic studies of *Trichodesmium* colonies obtained from the Atlantic and the Pacific oceans.

## Results and Discussion

A total of 52 metagenomic-assembled genomes (MAGs) were obtained from *Trichodesmium* colonies, of which 42 MAGs were of high quality (90% complete and <5% redundancy (Supplementary Table 1) (27). The relative abundance of the different MAGs was consistent across the three samples with *T. thiebautii* MAG 52 making up 71% ± 6 of the total relative abundance (Supplementary Table 1). In parallel to the metagenomic analysis, a proteomic dataset consisting of 14161 peptide sequences matching to protein sequences of 52 different MAGs further probed the activity within the Red Sea *Trichodesmium* consortia (Supplementary Table 2; Supplementary Table 3).

### *Trichodesmium* colonies host diverse and flexible associated bacteria

*Trichodesmium* colonies consisted of a single *Trichodesmium* species (*T. thiebautii* MAG 52) and a diverse consortium of 51 associated bacteria spanning at least 10 different taxonomic orders (Figure 1). Unlike a previous study (28), our samples did not include any non-diazotrophic *Trichodesmium* populations and the 51 MAGs hereby represent the associated bacteria of *T. thiebautii* MAG 52. The majority of the associated bacterial MAGs belonged to taxonomic orders known to be present in *Trichodesmium* colonies and included the orders Rhodobacterales, Pseudomonadales, Balneolales, Rhodospirillales, Flavobacterales, Enterobacterales, Rhizobiales, and Sphingomonadales (9, 11–13).

**Figure 1.**
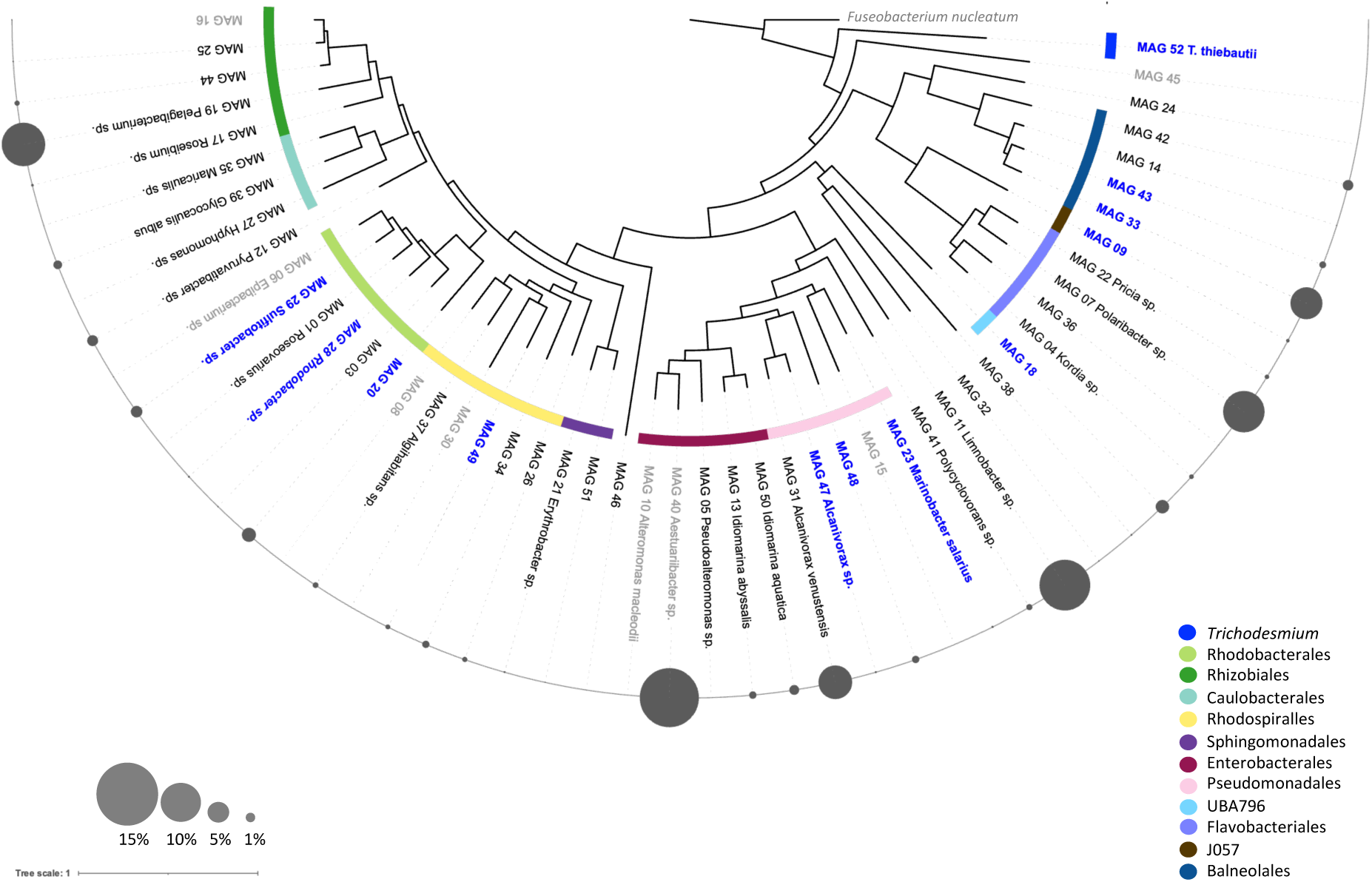
Phylogenetic tree of the 52 MAGs assembled from *Trichodesmium* colonies collected in the Red Sea. The tree is rooted by the outgroup *Fuseobacterium nucleatum*. Known bacterial orders are marked in different colors and several MAGs could taxonomically be identified at the genus level. The relative abundance of each MAG within the *Trichodesmium* consortium, excluding the abundance of *Trichodesmium* MAG 52 (±70% of samples), is presented as a bubble chart. The 11 MAGs that matched to MAGs assembled from a previous metagenomic dataset of Red Sea *Trichodesmium* colonies (28) are highlighted in blue and tentatively represent the core-consortium (see Supplementary Figure 2; Supplementary Table 4). MAGs that did not meet the requirements of a high-quality genome (>90% completeness; <5% redundancy) are indicated in grey. Note that re-occurring MAGs are not necessarily the most abundant MAGs present within the *Trichodesmium* colony. (a full taxonomic description of each MAG in this dataset can be found in Supplementary Table 1).

An intercomparison between the MAGs observed here and a previous metagenomic dataset from the Red Sea (28), identified a total of 11 MAGs present in both datasets (Supplementary Figure 2; Supplementary Table 4). These 11 MAGs included the particle-and algae-associated lineages (29–31) of *Marinobacter* sp. (MAG23) and *Alcanivorax* sp. (MAG 47) from the order Pseudomonadales; *Rhodobacter* sp. (MAG 28) and *Sulfitobacter* sp. (MAG29) from the order Rhodobacterales, and *Alteromonas macleodii* (MAG 10), a ubiquitous bacterium often found in association with phototrophs (32) and previously isolated from *T. erythraeum* IMS101 cultures (33). If these observations are consistent across a larger time-period of observations, these MAGs may represent a core component of the *Trichodesmium* consortium. Further comparison of our metagenomic dataset with those obtained from the Atlantic, North or South Pacific oceans, and the Red Sea (12, 13, 17, 18, 28), showed taxonomic similarities at the order level rather than at the species level. While most taxonomic orders re-occurred in all datasets, only one Balneolales genome (MAG 33) was present amongst all *Trichodesmium* metagenomic datasets. As none of the re-occurring MAGs represented highly abundant genomes within our samples (Figure 1), this may partially be the result of a higher sequencing depth (500 million reads) compared to past datasets. Nonetheless, the wide diversity coupled with the recurrence of only a few MAGs indicates the presence of a flexible assemblage of associated bacteria, which may have important implications for the community structure and functioning of the *Trichodesmium* consortium.

### Trichodesmium associated bacteria synthesize diverse photolabile siderophores, which may increase iron availability from dust

We explored the different Fe-uptake mechanisms present within the *Trichodesmium* consortium, as N_2_ and C fixation in *Trichodesmium* is often constrained by Fe availability (23, 34). Despite the high Fe-requirements of *Trichodesmium* (23)*, T. thiebautii* (MAG 52) only contained genes for the uptake of free Fe^+2^ and Fe^+3^, but not organic Fe, suggesting it is limited to a narrow range of forms it can directly take up (Figure 2). In contrast, MAGs of the associated bacteria contained a larger diversity of Fe-uptake genes which included the ability to take up organically bound Fe, such as heme, citrate, and siderophores (Figure 2). The proteome of the *Trichodesmium* consortium indicated that, at the time of sampling, *Trichodesmium* was actively taking up Fe through the presence of a ferric Fe-uptake transporters (FbpC, FbpA) and Fe-storage proteins (Dps, Bfr) (Supplementary Table 2).

**Figure 2.**
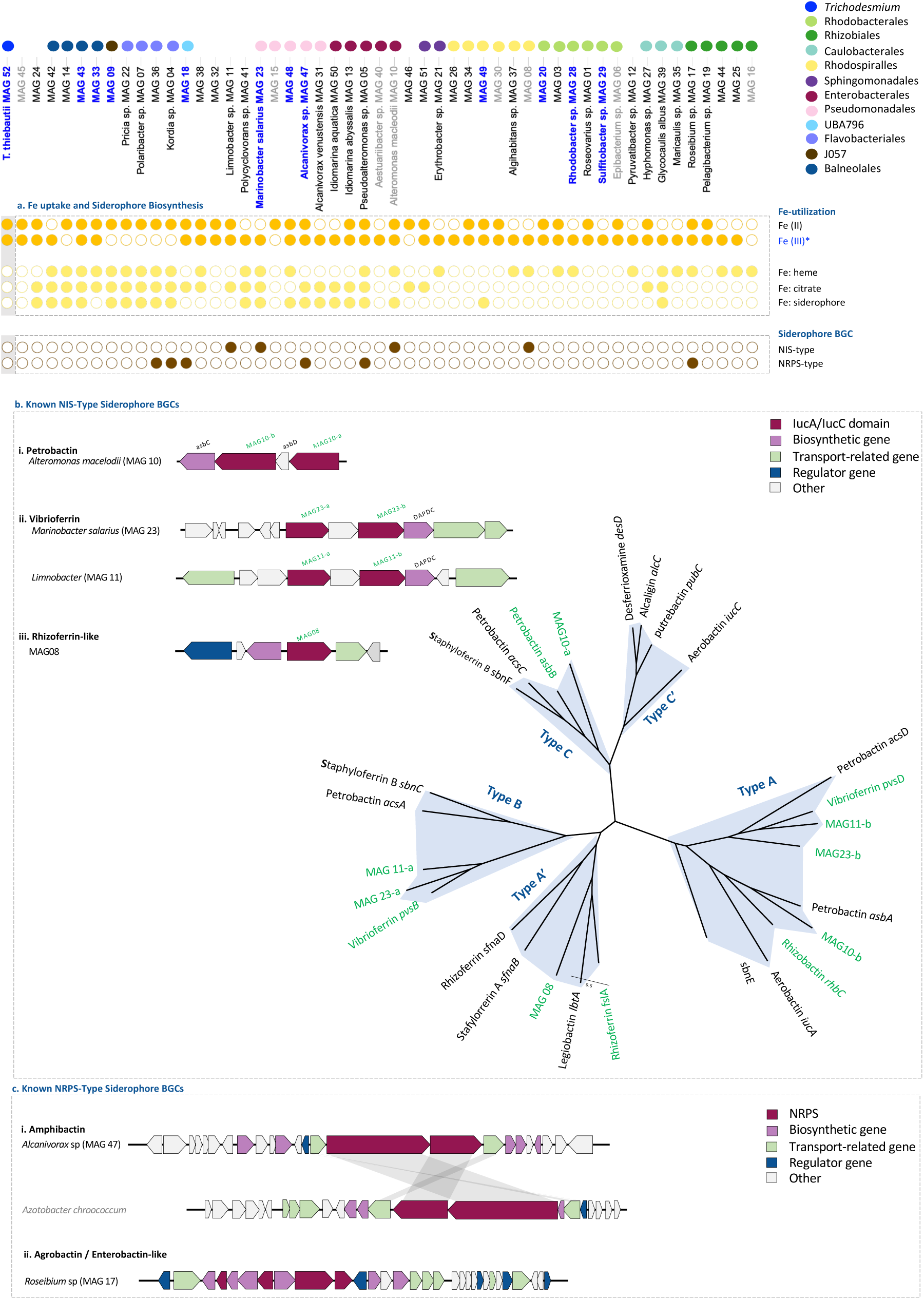
MAGs were probed for Fe-uptake and siderophore biosynthesis genes which are noted in Supplementary Table 5. **(a)** *T. thiebautii* MAG 52 does not contain a wide diversity of Fe-uptake genes, and the majority of MAGs of associated bacteria contain an uptake receptor for organic Fe (heme, citrate, siderophores). Several MAGs of associated bacteria contained an NRPS or NIS-type siderophore biosynthesis pathways (BGCs). **(b)** Siderophore biosynthetic gene clusters (BGCs) of known NIS-type siderophore BGCs can be clustered according to the presence of one or multiple iucA/iucC domains which enables one to target the closest siderophore BGC match (36). Using this method, we were able to link these pathways to the synthesis of the photolabile **(i)** petrobactin, **(ii)** vibrioferrin and **(iii)** rhizoferrin. (c) Siderophore BGCs of known NRPS siderophores. Using antiSMASH 7.0, which aligns BGCs to known siderophores within the MIBiG database, we were able to link 2 of the 6 putative siderophore NRPS-type pathways to **(i)** the membrane-bound amphibactin and **(ii)** agrobactin/enterobactin. (See Supplementary Figure 3 for more details on the siderophore biosynthesis pathways).

The genomes of 10 *Trichodesmium* associated bacteria spread across 6 different taxonomic orders encoded different siderophore biosynthesis gene clusters (BGCs) (Figure 2). The presence of a siderophore biosynthesis pathway was strain-specific and not present in an entire clade or lineage. Similar to previous studies, *T. thiebautii* MAG 52 did not contain any known siderophore biosynthesis pathways (12, 35). Siderophore BGCs could further be separated into 4 NIS-type and 6 NRPS-type siderophores (36, 37). Based on the presence of one or more *iucA/iucC* domains, the 4 NIS-types BGCs matched closely to petrobactin (*Alteromonas macleodii* MAG 10), vibrioferrin (*Marinobacter salarius* MAG 23; *Limnobacter sp.* MAG 11), and rhizoferrin (Rhodospirillales MAG 08) biosynthesis pathways (Figure 2). From the 6 different NRPS-type siderophores, only 2 matched a known BGC, namely that of the membrane-bound amphibactin (100%) (*Alcanivorax venustensis* MAG 47) and one resembling the BGC of the siderophores agrobactin (93%) or enterobactin (±83%) (*Roseibium* sp MAG 18) (Figure 2). Our results additionally indicated the presence of several novel NRPS-type siderophore biosynthesis pathways (Supplementary Figure 3). NRPS-type-siderophore BGCs are not as well characterized as NIS-type-siderophores BGCs and can be difficult to distinguish from other secondary metabolites such as toxins and antibiotics (38). While the 4 NRPS-type-siderophore BGCs could not be linked to known siderophore BGCs, several unknown metallophores have previously been identified and associated with *Trichodesmium* colonies and we speculate that some may be linked to these uncharacterized NRPS BGCs (19). The presence of multiple NRPS pathways containing a putative siderophore TonB-dependent receptor (SMCOG1082) likely involved in Fe uptake, highlights the potential for novel siderophore pathways within the *Trichodesmium* consortium and indicates the importance of additional culture-based research linking such pathways to a siderophore molecule.

Our results indicate that the *Trichodesmium* consortium in the Red Sea may interact with photolabile siderophore-producing bacteria to mine Fe from dust. The re-occurring consortium members *Marinobacter salarius* (MAG 23), *Alcanivorax venustensis* (MAG 47), and *Alteromonas macleodii* (MAG 10) are all able to synthesize the NIS-type siderophores petrobactin, vibrioferrin, and rhizoferrin (Figure 2) whose citrate-moiety results in their photolability (39). In the natural illumination of surface waters, where *Trichodesmium* resides, sunlight can reductively disassociate Fe from these siderophore complexes and provide a flux of highly available dissolved inorganic Fe (40, 41). While *Trichodesmium* does not contain a TonB-receptor, the presence of photolabile siderophores implies that Fe is also bioavailable to consortium members that lack a specific TonB-receptor for that particular siderophore. These photolabile siderophores act as a “common good” within the *Trichodesmium* consortia, where only a few members carrying the trait are needed to benefit the entire consortium (42). This contrasts with the uptake mode of photostable siderophores, such as desferrioxamine-B (DFOB), which require a specialized TonB-siderophore transport system (43). Similarly, amphibactin is a membrane-bound siderophore (44), which prevents its diffusion in the environment and subsequently its ability to act as a public good to the *Trichodesmium* consortium. A recent gene-knockout study in *Alteromonas macleodii* verified the ability of petrobactin to enhance the bioavailability of particulate Fe (22). This further supports the idea that *Trichodesmium* can interact with consortium members that produce photolabile siderophores to acquire Fe from dust.

Most of the siderophore BGCs detected here were identified in ubiquitous and copiotrophic bacteria typically found in association with particles and algae (11, 22), including *Marinobacter salarius* (MAG 23), *Alcanivorax venustensis* (MAG 47), and *Alteromonas macleodii* (MAG 10) (Figure 2). Siderophore-production within the *Trichodesmium* consortium appears to be linked to a lifestyle associated with a rapidly changing environment rich in N and C, where regulating Fe-bioavailability can provide a competitive advantage. In line with particle and algae associations, siderophore production has also been associated with other ecological functions, including heavy metal detoxification and intra-specific communication (45) (Supplementary Figure 4; Supplementary Table 5). Whether siderophore production can benefit the *Trichodesmium* consortium in functions other than Fe-dissolution from dust requires further investigation. Nonetheless, the presence of a few, yet reoccurring, siderophore producing MAGs augments previous measurements of siderophore production within the consortium (16, 19, 20) and reveals additional layers of *Trichodesmium* colonies’ intriguing ability to sequester Fe from dust.

### The *Trichodesmium* consortium is heterogeneous in genes related to the uptake and use of inorganic and organic phosphorous

We predicted that the majority of bacteria associated with *Trichodesmium* have the ability to metabolize phosphite or phosphonate. This can offer a key competitive advantage amongst consortium members in a P-limited environment (18, 46) including the Gulf of Aqaba in the Red Sea (47). Within a low-P environment, dissolved inorganic phosphorus is scarce, and bacteria must scavenge phosphate from dissolved organic phosphorus to meet their P-requirements (48). *Trichodesmium* is highly efficient at scavenging and utilizing organic-P through the production of alkaline phosphatase (AP), and the uptake of phosphonate and phosphite (49). Multiple studies have shown that *Trichodesmium* colonies in the Atlantic and Pacific Ocean are hotspots for the reduced P-cycling of phosphite or phosphonate (46, 50) suggesting that the majority of consortium members can process reduced-P.

Confirming past studies (18, 51), *T. thiebautii* MAG 52 encodes a set of functions to transport and metabolize diverse forms of P (Figure 3). This includes the presence of a high-affinity phosphate transporter (*pst*SCAB), and a phosphonate/phosphite transporter *(phn*CDE/*pxt*ABC). In addition to these transport systems, genes related to utilization and metabolism of organic-P included alkaline phosphatase (AP) (*pho*A*, pho*X), and a carbon-phosphorus (C-P) lyase (*phn*GHIJKLM) which hydrolyses a broad range of phosphonate compounds. *T. thiebautii* MAG 52 also encodes phosphite dehydrogenase (*ptxD)* which catalyzes the oxidation of phosphite to phosphate. The presence of PstB, PhoX, and PhnD within the proteome (Supplementary Table 2), suggests *Trichodesmium* was taking up phosphate, phosphonate, and hydrolyzing phosphoester compounds to meet its P demand. Several of these proteins are regulated by P (51), and their detection in the proteome is consistent with intense competition for P in the low P Red Sea.

**Figure 3.**
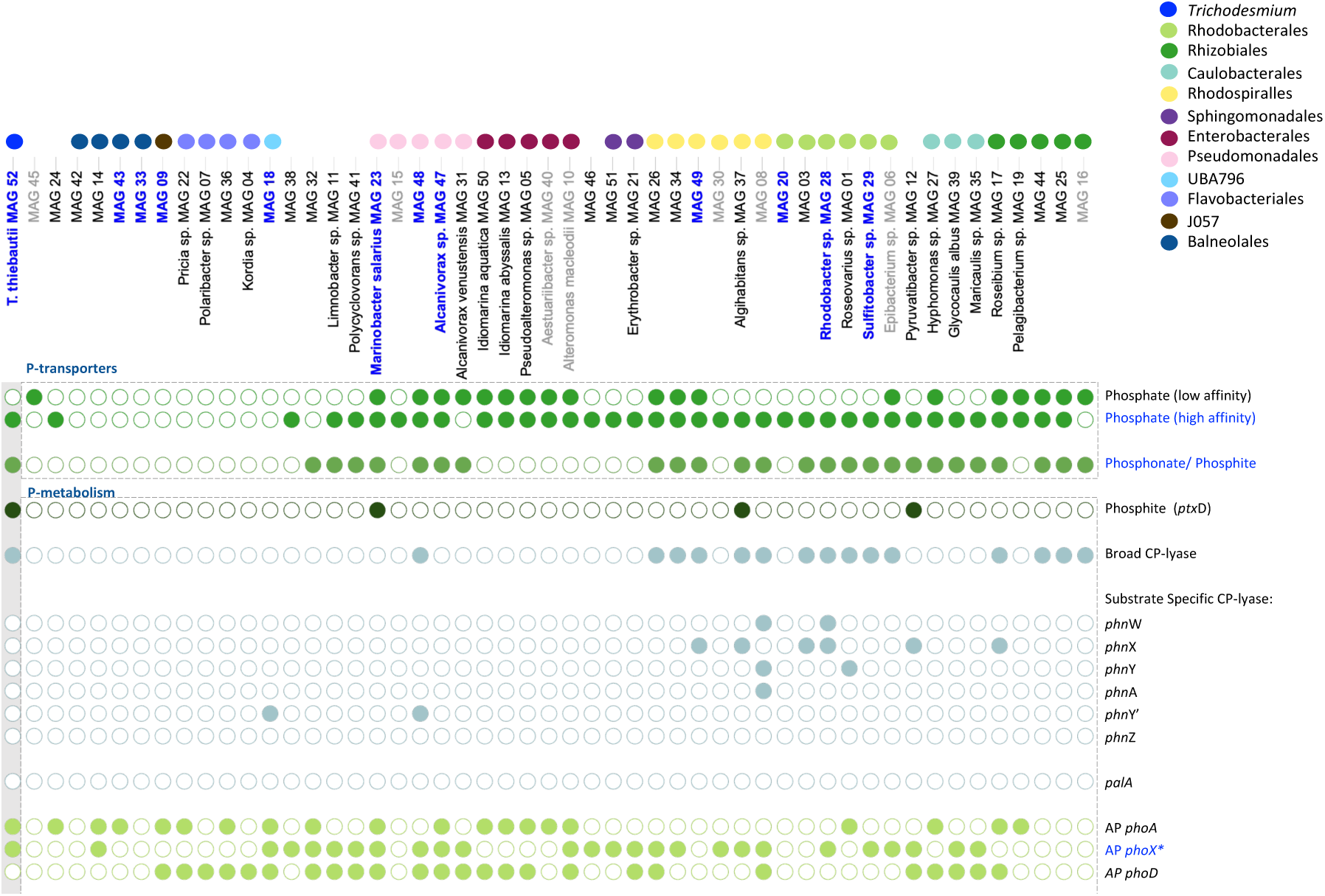
MAGs were probed for several different P-metabolism genes which are noted in Supplementary Table 5. *Trichodesmium* is known to be highly competitive for P, as exemplified by the multitude of **(a)** P-uptake systems for phosphate (*pst*S, *pst*B), phosphonate (*phn*CDE), and phosphonate/phosphite (*phn*CDE/*ptxABC*). **(b)** Within the *Trichodesmium* consortium, the ability to metabolize a range of phosphonate compounds using a broad specificity C-P lyase (*phn*GHIJKLM), is more prominent than the substrate-specific hydrolysis of 2-aminoethylphosphonate (2-AEP; *phn*WXYZA) or phosphonopyruvate (*pal*A) and phosphite metabolism (*ptx*D).

As KEGG annotation cannot accurately distinguish between a phosphonate (*phn*CDE) and a phosphite (*pxt*ABC) transporter, the presence of adjacent P-metabolism genes can further indicate the P-source linked to it. Specifically, the presence of phosphite dehydrogenase (*ptxD)* points to phosphite metabolism while a broad-specificity C-P lyase (*phnGHIJKLM*) or the substrate-specific hydrolysis of phosphonopyruvate (*pal*A) or 2-aminoethylphosphonate (2-AEP) to acetaldehyde (*phn*WX), acetate (*phn*WYA), or glycine (*phn*Y’Z), can point to the metabolism of phosphonate (52, 53). Our results showed that the presence of a *phn*CDE/*ptx*ABC uptake system was specific to certain taxonomic orders including Pseudomonadales, Rhodospirillales, Rhodobacterales, Caulobacterales, and Rhizobiales (Figure 3). Within the *Trichodesmium* consortium, these taxonomic orders may have a competitive advantage over members that lack them, particularly in low P conditions. Of these 25 MAGs, 16 could be linked to phosphonate metabolism through the presence of a broad-specificity C-P lyase. In contrast, the presence of phosphite dehydrogenase was species-specific where only 4 MAGs appeared to be able to process phosphite spread across different taxonomic orders (Figure 3). Within the *Trichodesmium* consortium, phosphonate metabolism appears to be the more common reduced P-currency of the two. 7 MAGs contained genes associated with the substrate-specific hydrolysis of 2-AEP but pathways were not complete, making it difficult to draw conclusions (Figure 3). Nonetheless, the presence of a broad-specificity C-P lyase appeared to be the most prevalent strategy for phosphonate utilization within the *Trichodesmium* consortium. This finding is in contrast to a meta-analysis of marine metagenomes where the most common phosphonate degradation strategy within the water column was the substrate-specific hydrolysis of 2-AEP (54). 2-AEP is the most abundant phosphonate source in the marine environment (55), and the differences in enzyme distribution within the Trichodesmium consortium may reflect heterogeneity in the phosphonate compounds present in the colony, relative to the water column. Differences in enzyme distribution may also be a function of the taxonomic composition of *Trichodesmium* associated bacteria relative to those in in the water column. Regardless, these observations suggest that a diversity of P compounds are being cycled by members of the consortia in the low P Red Sea and offer a new reference dataset for future comparisons with *Trichodesmium* consortia from other regions.

### Denitrification and dissimilatory nitrate reduction to ammonia (DNRA) pathways are modular in the *Trichodesmium* consortium

We explored the presence of multiple N-metabolic pathways as they are predicted to influence the biogeochemical contribution of *Trichodesmium* regarding N_2_ fixation. Our results showed that, in the *Trichodesmium* colonies sampled from the Red Sea, N_2_-fixation appears to be uniquely attributed to *T. thiebautii* MAG 52 (Figure 4). This contrasts with a previous study that has observed *nifH* genes in both *Trichodesmium* and their associated bacteria (12). In our study, however, only *T. thiebautii* (MAG 52) contained the full N2 fixation pathway (*nifDKHEB*: M00175) and therefore the ability to fix N2 within the colony. Similar to other studies (12, 17, 56), genes of the two additional N-transformation pathways of dissimilatory nitrate reduction to ammonium (DNRA) and denitrification were present within the *Trichodesmium* consortium (Figure 4). Respectively, denitrification and DNRA are N-loss and N-recycling processes and are predicted to enable the residing community to conserve and fully utilize the N2 fixed by *Trichodesmium* (12) but also alter the net contribution of *Trichodesmium* colonies to local N-cycling dynamics (15, 57). Our results indicated a relatively high number of 24 MAGs encoding genes involved in DNRA and 10 MAGs encoding genes involved in denitrification (Figure 4). Intriguingly, none of the MAGs analyzed contained the complete pathways for denitrification and DNRA, with the exception of Rhizobiales for the latter (Figure 4). Collectively, MAGs were able to complete all steps within either pathway, indicating that these pathways appear to be modular within the *Trichodesmium* consortium, where a bacteria can perform one step within the pathway but not necessarily all steps. While Rhizobiales (MAG 16, 17 and 44) contained the full DNRA pathway, which includes the reduction of nitrate to nitrite (*napAB, narGH*), and nitrite to ammonium (*nrfAH, nirBD*), the remaining MAGs contained only one of two steps (Figure 4). Similar to DNRA, several MAGs from the phyla Alphaproteobacteria contained a part of the denitrification steps, including the reduction of nitrite to nitric-oxide (*nir*KS) and nitric-oxide to nitrous oxide (*nor*BC), while members of the phyla Bacteroidetes could perform the final step within the denitrification pathway (*nosZ)* which converts nitrous oxide to nitrogen (Figure 4).

**Figure 4.**
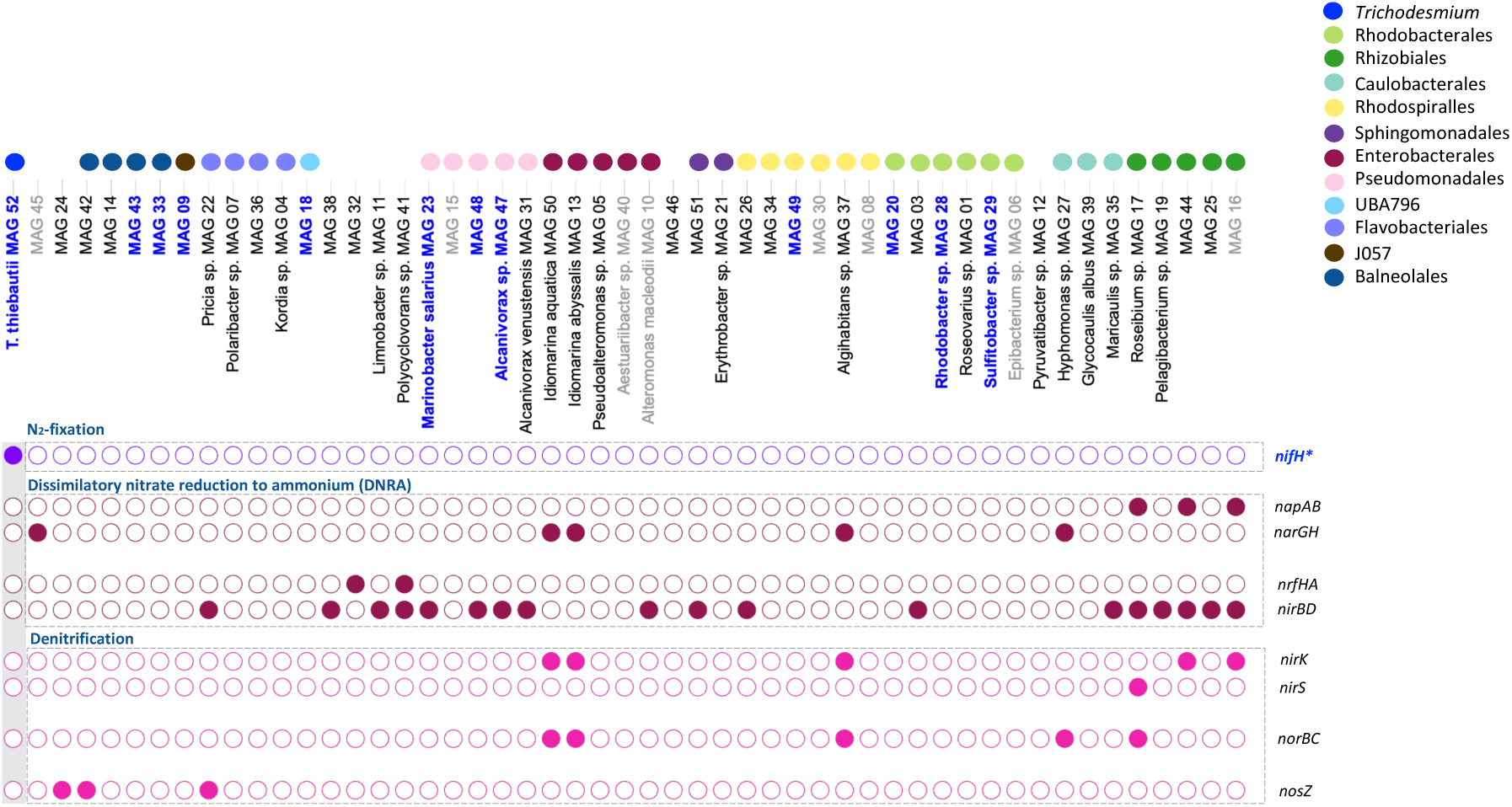
MAGs were probed for N-metabolic pathways and the genes for each step within the pathway are noted in Supplementary Table 5. *T. thiebautii* MAG 52 is the only MAG with a N_2_ fixation pathway. Several MAGs contained one or two steps within the pathways of denitrification or dissimilatory nitrate to reduction to ammonia (DNRA) indicating their modularity. Collectively, MAGs completed the denitrification and DNRA pathways, but none contained the genes for the entire pathway.

While the presence of modularity in denitrification and DNRA pathways itself is not uncommon (58–60) finding it present within the *Trichodesmium* consortium highlights the spatial role *Trichodesmium* plays in providing a platform where each step can be performed by a distinct subset of the microbial consortium. While the re-occurring consortium members *Alteromonas macleodii* (MAG 10)*, Marinobacter salarius* MAG 23), and *Alcanivorax sp* (MAG 47, MAG 48) encoded genes (*nir*BD) within the DNRA pathway, it did not appear to be a key characteristic within the *Trichodesmium* associated bacteria of the Red Sea. The presence of DNRA and denitrification pathways is rare in the oxic surface ocean, as these processes require anoxic and suboxic conditions (61). In the surface waters, suboxic as well as anoxic micro-environments were detected in marine aggregates, such as marine snow and fecal pellets, where enhanced microbial respiration results in the consumption of O_2_ (62, 63). *Trichodesmium* colonies may represent an additional environment where such processes may be taking place within the surface waters of the ocean. While it is not entirely clear when and where these N-cycling pathways are actively taking place within the colony studies have measured DNRA and denitrification processes within natural *Trichodesmium* colonies (15, 57). While the obligatory suboxic conditions for denitrification and DNRA may occur in the presence of dust and the rapid O_2_ consumption by bacteria (64), the formation of these microenvironments have thus far not been observed in *Trichodesmium* colonies (57). Altogether, these findings pinpoint the intricate community structure needed for such modular processes to take place within the consortium. The exact tradeoffs between *Trichodesmium* and denitrifying or DNRA bacteria, particularly in light of a changing ocean with higher dust deposition requires further investigation.

### *Trichodesmium* consortium members are interdependent for vitamins B1, B7 and B12

We explored the presence of vitamin auxotrophy within the consortium as an indicator for community interdependence. Previous studies have shown that *Trichodesmium* can synthesize and secrete the vitamin cobalamin (B12) to the environment (17). Our results show that while *Trichodesmium* MAG 52 encodes the biosynthesis pathways for B12, biotin (B7) and thiamin (B1), the majority of associated bacteria (49 of the 52 MAGs) lack a biosynthesis pathway for at least one of these vitamins (Figure 5). Bacteroidetes, Sphingomonadales, and Enterobacterales species, including *Alteromonas macleodii* (MAG 10), were all auxotrophic for vitamin B12. MAGs from the order Rhodospirillales lacked the ability to synthesize vitamin B7, whereas Rhizobiales and Rhodobacterales appeared to be auxotrophic for both vitamin B1 and B7. Only *Pseudoalteromonas* encoded all three biosynthesis pathways, including the re-occurring taxa *Alcanivorax* sp. (MAG 47) and *Marinobacter salarius* (MAG 23). These consortium members may also contribute to the synthesis and provision of vitamins B1, B7, and B12 to their auxotrophic counterparts (Figure 5) but, based on overall relative abundance, *Trichodesmium* represents the most likely source of B vitamins. In sum, the majority of associated bacteria (49 of 52 MAGs) were predicted to be auxotrophic for B12 and likely dependent on *Trichodesmium* to meet their intracellular requirements. Our results indicate that the production and secretion of different vitamins by *Trichodesmium* is a key determinant of community composition, stimulating vitamin-based interactions among auxotrophic consortium members.

**Figure 5.**
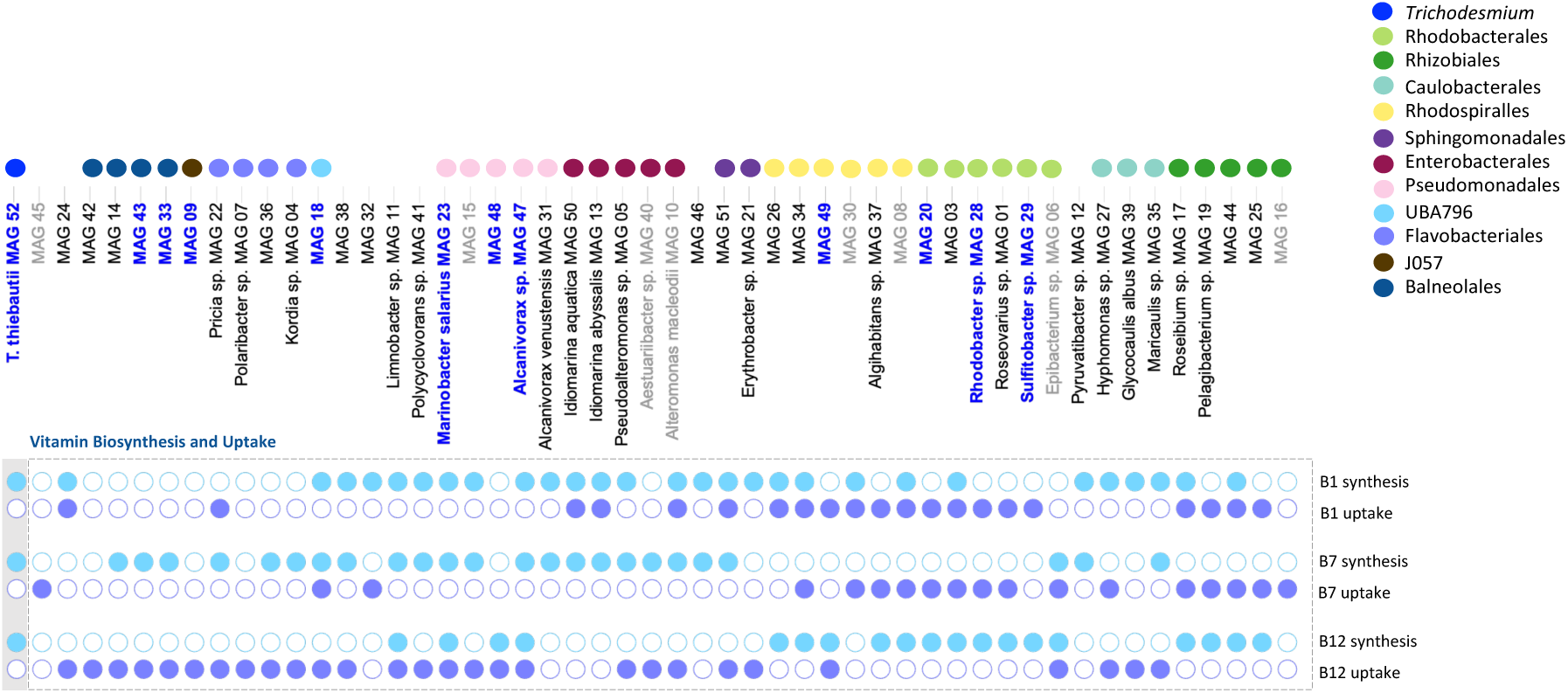
MAGs were probed for vitamin B1, B7, and B12 biosynthesis and uptake pathways. The genes used to indicate the biosynthesis or uptake of these vitamins are noted in Supplementary Table 5. *Trichodesmium* contains the biosynthesis pathways for all three vitamins but the vast majority of associated bacteria were auxotrophic for one or more vitamins. These results suggest that the associated bacteria rely on *Trichodesmium* to obtain their vitamin requirements.

### Interactions between copiotrophic microbes underpin nutrient cycling in *Trichodesmium* **consortia**

A literature study of the bacterial lineages present within the *Trichodesmium* consortium describes these taxonomic orders as typical primary colonizers of marine surfaces such as algae or sinking particles (29–31). Strains of *Alteromonas*, *Marinobacter,* and *Pseudomonas* often co-occur with phototrophs such as cyanobacteria as well as oil spills, likely as a result of their ability to degrade hydrocarbons and aromatic compounds (32, 65). While the prevalence of these copiotrophic lifestyles within the *Trichodesmium* consortium has been observed in previous studies (11, 13) it is still unclear whether their association with *Trichodesmium* is specific or reflective of a more general interaction between algae or particle associations. Copiotrophy is a widely encompassing term, but it generally concerns opportunistic bacteria with a versatile metabolic capacity to use a wide range of substrates, acting as important remineralizers by rapidly responding towards areas enriched in organic matter (66, 67). It is therefore possible to consider the majority of the consortium as active remineralizers of the C, N, P, Fe, and vitamins *Trichodesmium* is providing. Viewing *Trichodesmium* as a hotspot for nutrients is supported by our proteomic samples where the majority of protein hits attributed to *T. thiebautii* MAG 52 were related to photosynthetic (K05376; K05377; K02284) and diazotrophic machinery (K03839, K02588, K02591) indicating that the processes of N and C fixation were actively taking place at the time of collection (Supplementary Table 2). In turn, the majority of protein hits associated to consortia members were involved in the uptake of amino-acids (K01999; K09969), sugars (K02055; K02027; K10552), and peptides (K02035; K02032) which is reflective of typical copiotrophic activity (Supplementary Table 3).

Copiotrophic bacteria are also characterised by their strong ability to interact within their associations (68). This is supported by the prominence of several bacterial-interactive and particle-associated traits in the MAGs studied here (68, 69) (Supplementary Figure 4; Supplementary Table 5). The presence of bacterial-associated interactions taking place between *Trichodesmium,* and their consortium members could further be implied through the presence of several proteins involved in pilus formation (K02658, K02662, K02669) and secretion systems (K11003; K03072; K03070, K03110, K03073; K03217) within *T. thiebautii* MAG 52 (Supplementary Table 2). Collectively, these findings are indicative of an intricate network of interaction between *Trichodesmium* and its associated bacteria. For example, *T. thiebautii* MAG 52 and several MAGs, including re-occurring members from the order Rhodobacterales, contained a homoserine lactone biosynthesis pathway (Supplementary Figure 4a). Previous studies have predicted that consortium members can directly interact with *Trichodesmium* through the production of homoserine lactones (HLs), where the presence of

HLs was shown to modulate N_2_ fixation rates (14) and increase alkaline phosphatase (AP) activity in *Trichodesmium* (70). The functional association of poorly studied bacterial clades, in particular Balneolales (Supplementary Table 1), however, remained more elusive and indicated that we still lack a clear understanding of these bacteria and their role within the *Trichodesmium* consortium. Further investigations will be required to untangle the different dynamics of both general and specific interactions taking place between members of the *Trichodesmium* consortium.

### Conclusions and Future Perspectives

The complex *Trichodesmium* consortium represents an ideal system to study the intricate interactions taking place within a particle-rich system (Figure 6), asking how these processes may impact the C, N, P and Fe-cycles in a rapidly changing ocean where both microbial community composition and changes in microbial abundance are expected to occur (25, 71). The *Trichodesmium* consortium is redundant for multiple traits related to nutrient cycling including siderophore production, denitrification, DNRA, and reduced phosphate uptake. The processes of denitrification and DNRA were modular, indicating an interactive and spatial coupling between different consortium members. The associated bacteria were largely auxotrophic for one or more vitamins that *Trichodesmium* can synthesize, and this is likely a major trait influencing the community structure (Figure 6). The vast majority of the associated bacteria could further be characterized according to a copiotrophic lifestyle where *Trichodesmium* represents a hotspot for C, N, P, Fe, and vitamins to the residing community. On the other hand, Fe-uptake through the production of siderophores appeared to be a key beneficial function *Trichodesmium* can obtain from its residing community. *Trichodesmium* harbored several bacteria that can secrete photolabile siderophores, supporting the idea that *Trichodesmium* can meet their high Fe-demand by interacting with siderophore-producing bacteria to mine Fe from dust particles collected within the colony. Other, as of yet uncharacterized, traits will likely further elucidate the factors that drive the community structure of *Trichodesmium* colonies. Our results highlight the presence of a dynamic and flexible consortium where functional traits are well conserved within the *Trichodesmium* consortium, underpinning the resilience of the colony within an ever-changing ocean.

**Figure 6.**
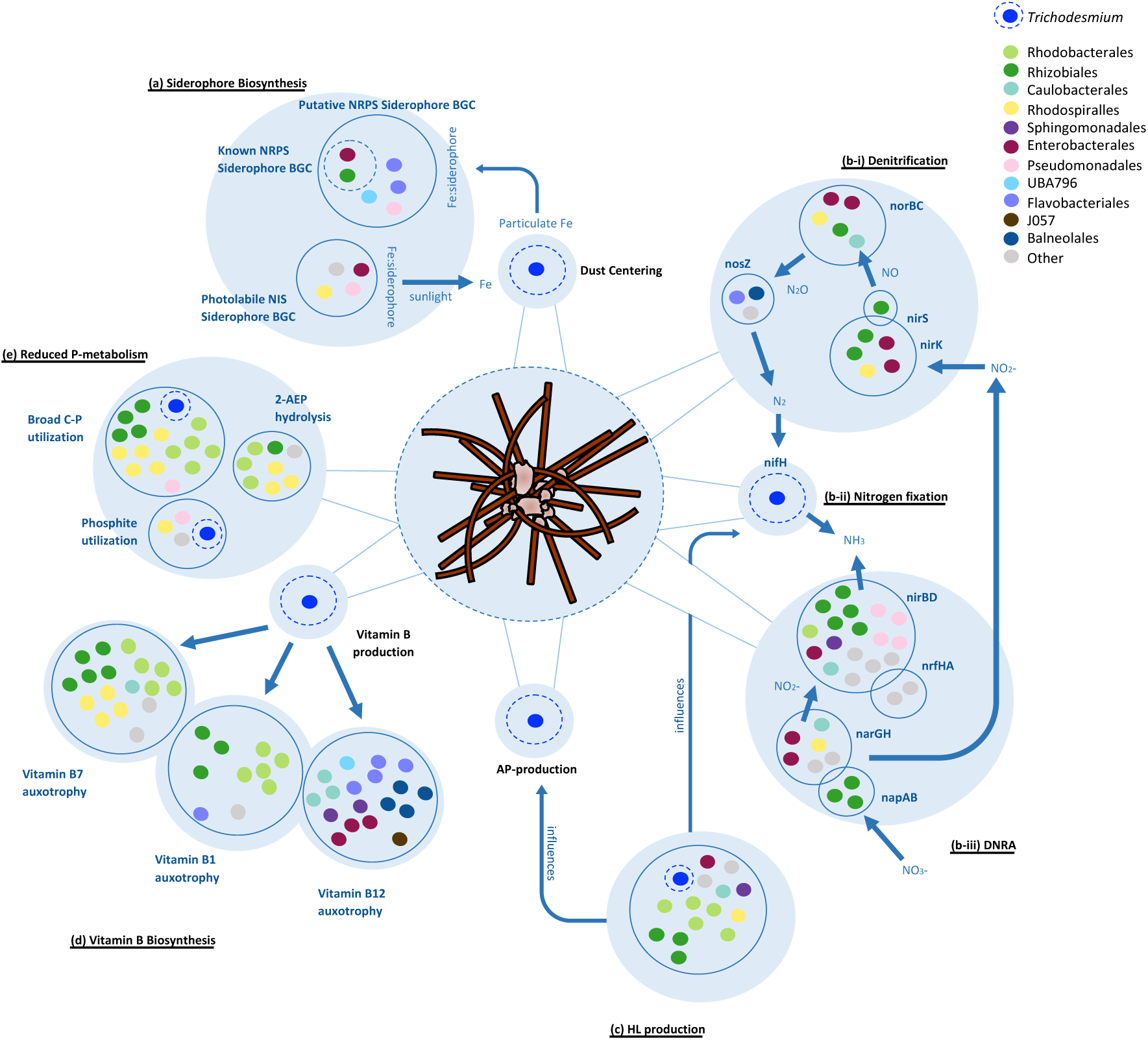
Schematic summary of the potential interactions taking place within the Trichodesmium consortia as discussed in this study. Genes for each trait are noted in Supplementary Table 5. Each blue bubble indicates a different function, and each blue circle pools together the different MAGs containing a particular trait for that function. Each MAG is represented by a colored dot reflective of the taxonomic order it is associated with (grey dots represent MAGs that were not attributed to a specific taxonomic order). *Trichodesmium* is highlighted in cobalt-blue, and the important functions *Trichodesmium* is known to display are highlighted with a dotted circle. (See Supplementary Table 5 for the full list of functional traits and their presence/absence). **(a)** Several siderophore biosynthesis genes are present in a diverse set of MAGs. Siderophores are predicted to mine Fe from dust that has been collected and centered within *Trichodesmium* colonies. 6 of the 4 were identified as known NIS-type siderophores of which petrobactin, vibrioferrin and rhizoferrin are photolabile. In the presence of sunlight, the Fe complexed to these siderophores disassociates and acts as a public good to the larger consortium. The genes and pathways for **(b-i)** denitrification, **(b-ii)** N_2_ fixation, and **(b-iii)** dissimilatory-nitrate-reduction-to-nitrate (DNRA) are present in several MAGs of the consortium. These processes appear to be modular within the *Trichodesmium* consortium as different taxonomic groups complete the steps for these pathways. **(c)** Several MAGs contain a homoserine lactone synthesis pathway (HL), including *Trichodesmium*. HL acts as an intercommunicating quorum sensing molecule and has been shown to influence the production of alkaline phosphatase (AP) and N_2_ fixation in *Trichodesmium*. (associated bacteria can also produce AP but is not depicted here - see Figure 3). **(d)** *Trichodesmium* is capable of producing the vitamins cobalamin (B12), thiamin (B1) and biotin (B7). Different subsets of MAGs lack the biosynthesis pathways for one or more of these vitamins with very few capable of synthesizing all three (associated bacteria can also produce vitamins but is not depicted here - see Figure 5). Consortia members and *Trichodesmium* likely exhibit an interdependent relationship for these vitamins. **(e)** *Trichodesmium* colonies have been shown to take up and utilize phosphonate and phosphite in a low-P environment and MAGs containing these pathways may have a competitive advantage over members that lack them.

## Material and Methods

### Trichodesmium metagenomic sampling and extraction

*Trichodesmium* puff colonies were hand-picked during the spring bloom (6^th^ May, 2019) and separated into three samples for metagenomics (∼100-200 colonies each) using a 100 µm phytoplankton net at 20 m depth in the Gulf of Aqaba (Eilat, Israel) (29.56°N, 34.95°E). Colonies were washed 3 times, by gently picking colonies with a pipette and placing them in new petri dishes with fresh 0.2 µm filtered seawater, before being filtered on a 0.2 µm, 45 mm polycarbonate (PC) filter using a vacuum pump. Filters were flash frozen in liquid N_2_ and kept at −80°C. DNA was extracted from the three *Trichodesmium* samples using the DNeasy Plant Pro Kit (Qiagen), with a minor modification to the lysis procedure. The kit-provided tissue disruption tubes were not used. Rather, ∼250 μL zirconia/silica beads (0.5 mm) were added to each sample tube before the addition of Solution CD1, and then samples were vortexed for 5 min. The resulting lysate was processed as per the remainder of the manufacturer instructions. DNA was quantified fluorometrically using the dsDNA HS kit (ThermoFisher Scientific) on a Qubit. From the three samples, metagenomic libraries were prepared and sequenced with 0.7 billion reads 2 × 151 bp reads on Illumina NovaSeq S4 at the DOE Joint Genome Institute (California, USA). This study can be found under JGI Gold Study ID number Gs0149370.

### Metagenomic raw read assembly and binning

Sequences were analysed from this study and compared to previously published metagenomic studies from the Red Sea (PRJNA804487) (28) and metagenomes of *Trichodesmium* colonies isolated from the Pacific (PRJNA435427) (17) (PRJNA358796) (12) and the Atlantic (PRJNA330990) (13) Oceans. All raw sequences were analyzed using the same protocol described below.

Metagenomes were assembled, binned, and quantified using the ATLAS (v2) pipeline (72). Briefly, raw sequences underwent quality control through the BBTools suite (73, 74) and were assembled using metaSPAdes (75) (k-mer lengths: 21, 33, 55, 99, 121 bp). MAGs were binned from each sample using MetaBAT 2 (76), MaxBin 2.0 (77) and VAMB (78). The completeness and redundancy of each bin were subsequently assessed using CheckM2 v0.1.2 (79). MAGs were considered of high-quality if they reached 90% completeness and 10% redundancy and were highlighted throughout our results and figures (27). A non-redundant set of bins was produced using DAS Tool (80) and dRep (81) based on an average nucleotide identity (ANI) cutoff of 97.5%. Genes were predicted using Prodigal (82). The coverage of each MAG was quantified across each sample by mapping reads back to each MAG. We refer to the MAGs from this study as MAG XX, bacterial MAGs from a previous Red Sea study as MAG R-XX (28) and bacterial MAGs from other metagenomic studies from the Pacific and Atlantic Oceans (12, 13, 17) as MAG T-XX.

### Phylogenetic Diversity

Metagenomically assembled genomes (MAGs) were taxonomically characterized using the genome taxonomy database tool kit GTDB-tk v2.1.1 (83). MAGs were classified according to GTDB taxonomy and NCBI taxonomy (Supplementary Table 1; Supplementary Table 4). GTDB taxonomy is used throughout the analysis, but to simplify comparisons with the literature, we used the larger NCBI taxonomic order Rhodospirillales which encompasses the smaller GTDB taxonomic groups of Kiloniellales and Thalassobaculaceae.

Phylogeny was inferred using GTo-Tree (v.1.16.12; default settings) (84) from a concatenation of 74 conserved single-copy HMM markers for bacteria using the best-fit model Q.pfam+R7 model in IQ-Tree (v2.1.3), using the Bayesian information criterion (BIC) (85, 86). Shimodaira–Hasegawa approximate likelihood-ratio test (SH-aLRT) and ultrafast bootstrap approximation (UFBoot) branch support values were estimated from 1000 bootstraps. The tree was visualized using FigTree (v1.4.4) and rooted using the genome of *Fuseobacterium nucleatum* (PRJNA1419) as an outgroup according to (87). MAG were considered to re-occur in another metagenomic dataset if its genome sequence identity was >97.5% similar (Supplementary Figure 1).

### Functional Diversity

The amino-acid sequence for each MAG was annotated using hmmscan and GhostKoala under default parameters (50% identity cutoff), utilizing the PFAM and KEGG databases, respectively (88, 89). Secondary metabolites (e.g. biosynthesis of siderophores and homoserine lactones) were screened for each MAG using AntiSMASH 7.0. In certain cases, the Rapid Annotation using Subsystem Technology (RAST) server was used to annotate amino-acid sequences using the collection of protein families, FIGfams (90, 91). In addition, each MAG was assessed using METABOLIC (v4.0) which lists the presence of interactive KEGG modules for each MAG using a default completeness cutoff of 0.75 (92). Based on these different annotation formats, we inspected each MAG for the presence of functional traits. For each functional pathway or gene, the annotation method and coinciding KEGG and PFAM ID number can be found in Supplementary Table 5. The presence and absence of different traits for each MAG was displayed using the Interactive Tree Of Life (iTOL) visualization tool (93).

### Iron metabolism

To explore the relationship between *Trichodesmium* and its associated bacteria for the bioavailability of Fe from dust, we searched MAGs for the KEGG IDs of multiple Fe-uptake pathways. Fe-uptake receptors were separated according to abiotic Fe^+2^ (*feoB*:K04759), Fe^+3^ (*afuA*:K02012), and organic Fe uptake which is further separated into Fe-heme (*hemR*:K16087), Fe-citrate (*fecA*:K16091) and Fe-siderophores (*fevS*:K02016). The presence of an Fe:siderophore receptor K02016 should be taken cautiously as its KEGG annotation does not clearly distinguish between siderophores or cobalamin as a substrate.

Siderophore biosynthesis pathways within MAGs were identified using a conservative approach that required a combination of both KEGG and HMM annotation for each MAG and the use of AntiSMASH 7.0 and FeGenie tools (94) (Supplementary Figure 4). Pathways were separately analyzed for both Non-Ribosomal Peptide Synthetase (NRPS) Type and NRPS Independent Synthetase (NIS) Type biosynthesis pathways. NIS-type siderophore biosynthesis pathways are more easily identifiable of the two, and we searched for the presence of the Fe-uptake chelate (*iuc)*A*/iuc*C domain (PF04183). NIS genes were clustered into different types using known NIS biosynthesis genes as defined and described by Carroll and Moore (36). Briefly, genes were aligned using the online Multiple Alignment Fast Fourier Transformation (MAFFT v7.490; L-INS-i) software (95). A phylogenetic tree was constructed from this alignment using IQtree2 (v2.1.3). The resulting best-fit substitution model Q.pfam+F+R4 was selected using Bayesian information criteria (BIC). Branches were assigned using Shimodira– Hasegawa approximate likelihood-ratio test (SH-aLRT) and ultrafast bootstrap approximation (UFBoot) branch support values were estimated from 1,000 bootstraps. The resulting consensus tree was visualized using FigTree (v1.4.4).

The identification of NRPS-type siderophore biosynthesis pathways required FeGenie, which utilizes biosynthetic pHMMs, and was used to highlight putative NRPS-type siderophores that were further inspected using AntiSMASH 7.0. MAGs containing an NRPS pathway coupled with a siderophore transport-receptor (SMCOG1082) within the gene-cluster were marked as putative NRPS-siderophore producers. These pathways were further compared to known NRPS siderophore pathways using the MiBIG database (96) and the KEGG pathway map for NRPS-type siderophores biosynthesis gene cluster (BGC) map01053.

### Vitamin B1, B7, and B12 biosynthesis and uptake

We assume that a bacterial strain is auxotrophic for thiamin (B1), biotin (B7) or cobalamin (B12) if their coinciding genome lacks a cobalamin biosynthesis pathway and contains a transporter for its uptake. We searched MAGs for the KEGG IDs and determined the presence of a vitamin biosynthesis pathway using a KEGG map for each vitamin. The presence of a thiamin biosynthesis pathway (M00127) was determined by the presence of *thi*DCEGL (K00946; K00788; K03149; K00941; K03147). A thiamin uptake system was present if a coinciding transporter system (M00192) was observed. The presence of a biotin biosynthesis pathway (M00581) was determined by the presence of *bio*ABDFH (K01012; K01935; K00833; K00652; K02170). A biotin uptake system was present if a coinciding transporter system (M00581) was observed. The presence of a cobalamin biosynthesis pathway was determined by the presence of *cob*ACDUQS (K19221; K02226; K02227; K02231:K02233). A putative vitamin B12 uptake system was considered present in a MAG if the specific transporter *btu*B (K16092) was observed.

### Nitrogen metabolism

N-metabolism pathways om MAGs were compiled from a selected set of KEGG ID numbers using METABOLIC-C (v4.0) and the KEGG pathway map ko00910. MAGs were queried for the presence of genes related to N_2_ fixation, dissimilatory nitrate reduction to ammonium (DNRA), and denitrification. The ability to fix N_2_ was determined through the presence of *nif*HKD (K02588, K02591, K02586). DNRA can be separated into two steps and includes the reduction of nitrate to nitrite (*nap*AB: K02567, K02568*; nar*GH: K00370-K00374) and nitrite to ammonia (*nrf*AH: K15876, K03385*, nir*BD: K00362, K00363). Denitrification can further be separated into the reduction of nitrite to nitric oxide (*nir*S:K15864*; nir*K: K00368), nitric oxide to nitrous oxide (*nor*BC: K04561, K02305) and nitrous oxide to N_2_ (*nos*Z: K00376).

### Phosphorus metabolism

The presence of multiple P-metabolic pathways within the *Trichodesmium* consortium were explored. We explored several transporters for high-affinity phosphate (*pst*SCAB: K02036-K02038, K02040), low-affinity phosphate (*pitA*:K03306), and a phosphite/phosphonate transporter (*phn*CDE: K02041; K02042; K02044). We searched MAGs for KEGG ID numbers for the presence of phosphite dehydrogenase (*ptx*D:K18916), which oxidizes phosphite to phosphate, and the broad specificity C-P lyase (*phn*GHIJKLM: K06162-6, K05780-1), which hydrolyses phosphonate bonds in organic P. Additionally, MAGs were queried for the presence of the substrate-specific phosphonate metabolism enzymes including *phn*A (K19670), *phn*X (K05306), and *phn*W (K03430) and collectively these genes were used to indicate the functional potential for reduced P-metabolism. Lastly, MAGs were examined for the presence of alkaline phosphatases (AP) *pho*A (K01077), *pho*X (K07093), and *pho*D (K01113). AP hydrolyses phosphoesters enabling the metabolism of organic-P.

### Bacterial and Particle Interaction traits

Known traits that are indicative of putative bacterial interactions were previously summarized and identified by Zoccarato *et al* for marine bacterial genomes (68). In addition to siderophore biosynthesis and vitamin B1, B7 and B12 biosynthesis (see previously), MAGs were probed for the presence of homoserine lactone (HLs) biosynthesis genes using AntiSMASH 7.0. The presence of auxin efflux genes was determined according to RAST annotation. MAGs were screened for KEGG ID numbers for the presence of the quorum sensing regulation gene *lux*R (K07782) and *bja*R1 (K18098), secretion systems (KEGG map03070), and motility and adhesion genes (KEGG map02020) (Supplementary Table 5). Traits involved in particle-interaction were previously described and summarized by Debeljak *et al* (69). In addition to Fe-uptake and siderophore biosynthesis pathways, MAGs were screened for KEGG ID numbers involved in metal-response, metal-efflux and metal-storage pathways (Supplementary Table 5).

### Trichodesmium proteomic sampling and extraction

In parallel to the metagenomic sampling, 20 puff colonies were picked, in triplicate, for proteomic analysis at the Environmental Molecular Sciences Laboratory (EMSL) at Pacific Northwest National Laboratory according to standardized protocols. Colonies were washed 3 times with filtered seawater, as described above. Samples were centrifuged, and the resulting pellet was diluted in 200 µl 8 M urea and transferred to 2 mL pre-filled Micro-Organism Lysing Mix glass bead tubes and bead beat in a Bead Ruptor Elite bead mill homogenizer (OMNI International, Kennesaw Georgia) at speed 5.5 for 45 sec. After bead beating, the lysate was transferred to a new 4 mL tube and immediately placed in an ice block and spun at 1,000 x g for 10 mins at 4°C. 200 µl of the lysed samples were transferred into 2 mL centrifuge tubes. A bicinchoninic acid (BCA) assay (Thermo Scientific, MA USA) was performed to determine protein concentration. Following the assay, 10 mM dithiothreitol (DTT) was added to the samples and incubated at 60°C for 30 mins with constant shaking at 800 rpm. Samples were diluted 8-fold for preparation for digestion with 100 mM NH_4_HCO_3_, 1 mM CaCl_2_ and sequencing grade trypsin (Promega, WI) was added to all protein samples at a 1:50 (w/w) trypsin-to-protein ratio for 3 hrs at 37°C, 450 rpm. Digested samples were desalted using a 4-probe positive pressure Gilson GX-274 ASPEC™ system (Gilson Inc., WI) with Discovery C18 50 mg/1 mL solid phase extraction tubes (Supelco, MO), using the following protocol: 3 mL of methanol was added for conditioning followed by 3 mL of 0.1% trifluoroacetic acid (TFA) in H_2_O. The samples were then loaded onto each column followed by 4 mL of 95:5: H_2_O:ACN, 0.1% TFA. Samples were eluted with 1mL 80:20 ACN:H_2_O, 0.1% TFA. The samples were concentrated using a speed vac and a final BCA was performed to determine the peptide concentration and samples were fractionated into 12 fractions for LC-MS/MS analysis.

MS analysis was performed using a Q-Exactive Plus mass spectrometer (Thermo Scientific) outfitted with a homemade nano-electrospray ionization interface. Electrospray emitters used 150 μm o.d. × 20 μm i.d. chemically etched fused silica (126). The ion transfer tube temperature and spray voltage were 300°C and 2.2 kV, respectively. Data was collected for 120 min following a 10 min delay after completion of sample trapping and start of gradient. FT-MS spectra were acquired from 300 to 1800 m/z at a resolution of 70 k (AGC target 3e6) and the top 12 FT-HCD-MS/MS spectra were acquired in data-dependent mode with an isolation window of 1.5 m/z at a resolution of 17.5 k (AGC target 1e5) using a normalized collision energy of 30, dynamic exclusion time of 30 s, and detected charge state of an ion 2 or higher. The resulting proteomic samples from this study was deposited in MassIVE (MSV000091416).

### Proteomic Analysis

Peptide matching was performed using MSGF+ software (97) against the amino-acid sequences obtained from the matching metagenome dataset through PROKKA (98) using the ATLAS pipeline (v2) (72) resulting in 201374 protein sequences. MSGF+ was used in target/decoy mode with 20 ppm parent ion tolerance, partial tryptic rule, and M-oxidation (+15.9949) as a dynamic modification. Best matches from the MSGF+ searches were filtered at 1% FDR based on the target/decoy model; this set of peptides was used in consequent quantitation analysis. Spectral counts were performed without consideration of peptide specificity for proteins and MAGs. 92% of peptides mapped to unique proteins while 93% of peptides were found in only one MAG. Spectral count normalization was performed using the normalized spectral abundance factor (NSAF) approach (99). Briefly, counts were divided by the matching peptide length and their relative abundance was calculated from the total number of spectral hits per sample.

## Acknowledgements

This research was performed under the Facilities Integrating Collaborations for User Science (FICUS) program (proposal: 10.46936/fics.proj.2018.50403/60000053) and used resources at the DOE Joint Genome Institute and the Environmental Molecular Sciences Laboratory, which are DOE Office of Science User Facilities. Both facilities are sponsored by the Biological and Environmental Research program and operated under Contract Nos. DE-AC02-05CH11231 (JGI) and DE-AC05-76RL01830 (EMSL). This research was supported by Grant no. 2020041 awarded to YS and RB from the United States-Israel Binational Science Foundation (BSF).

This research was supported by Grant no. 2020041 awarded to YS and RB from the United States-Israel Binational Science Foundation (BSF) and the Israel Ministry of Science and Technology (MOST) Grant no. 001126 to MR-B. Additional support was provided by a grant from the Simons Foundation award ID 721225 to STD.

## Data Availability Statement

Raw sequences are available under the Gold Study ID number Gs0149370. The 52 metagenomic assembled genomes analysed in this study are available at NCBI under the BioProject number PRJNA944101. Proteomic samples are available at MassIVE under the project accession number MSV000091416.

## Author Contributions

YS, RB, MG, SB, conceived of this study. SB, FZ, SW collected the DNA and protein samples. SB performed the experiments together with FZ, SW and YS. SD, SH, TG extracted and sequenced the DNA. CDN extracted and processed protein samples that were analyzed by NT and RB. MRB and CK performed the metagenomic analysis. CK wrote the manuscript under guidance of YS, MRB, RB, MG, SD, with further input from SW, SB, FZ, NT.

## SUPPLEMENTARY TABLES

**Supplementary Table 1.** General information about the MAGs making up the Red Sea *Trichodesmium* consortium. The relative abundance of associated bacteria (%) and standard deviation of the triplicates (SD) excludes the abundance of *T. thiebautii* (MAG 52). The relative abundance of MAGs was consistent across our samples. MAGs that did not meet the high-quality standard of 90% completeness and <5% redundancy are marked in green (10 of 52) and the functional analysis from these MAGs needs to be taken with more caution. The % of the genomes annotated by RAST is also indicated, showcasing how MAGs from the phyla Bacteroidetes are marked with a low annotation percentage. The closest taxonomic identity is given using both NCBI and GTDB-taxonomy. Throughout this paper we use GTDB-taxonomy except for the order Rhodospirillales.

**Supplementary Table 2:** Top proteomic hits matching to *T. thiebautii* MAG52. Spectrum counts that mapped to KEGG ID numbers mentioned in Supplementary Table 5 are highlighted in blue. Most proteins are geared towards the processes of photosynthesis, nitrogen fixation, carbon metabolism, amino acid uptake, protein synthesis, and phosphate uptake. Total spectral counts (T-total) and their associated KEGG or PFAM ID number and their description are presented together with the average relative contribution (T0%) of each protein hit to all Trichodesmium-related proteins and for all three samples (%T1-3). Collectively, the top 50 proteins represented >50% of the total spectral counts.

**Supplementary Table 3:** Top proteomic hits matching to MAGs of *Trichodesmium* associated bacteria. Many proteins are geared towards a typical heterotrophic and copiotrophic lifestyle including proteins related to amino-acid uptake and carbon metabolism are highlighted in bold. Total spectral counts (%) and their associated KEGG or PFAM ID numbers and their description are presented together with the average relative contribution (Avg%) of each protein hit to all Trichodesmium-related proteins and for all three samples (%T1-3).

**Supplementary Table 4.** **Taxonomic identity of MAGs used in this study are compared to MAGs assembled from other metagenomic studies of *Trichodesmium* colonies** (Red Sea samples: MAG R-XX (28), the Pacific (12, 17) and Atlantic samples (13): MAG T-XX. Their closest taxonomic identity is given using both NCBI and GTDB-taxonomy. Throughout this paper we use GTDB-taxonomy except for the order Rhodospirillales. Highlighted in blue are the 11 MAGs from this dataset that matched to those assembled from these other datasets (in bold), using a 97.5 % identity cutoff. The phylogenetic tree of all MAGs from all datasets combined can be seen in Supplementary Figure 2.

**Supplementary Table 5.** The presence and absence of functional traits separated into larger groups including Fe-uptake, P-metabolism, N-metabolism, Vitamin B synthesis and uptake, metal regulators, metal efflux, metal storage, signaling, secretion systems. motility and adhesion genes. The associated analysis tool, KEGG id number or module is given for each trait. The total number of MAGs containing a specific trait and (MAGs %) is calculated and show below. Traits that were detected within our proteomic samples are also highlighted in blue.

## SUPPLEMENTARY FIGURES

**Supplementary Figure 1.**
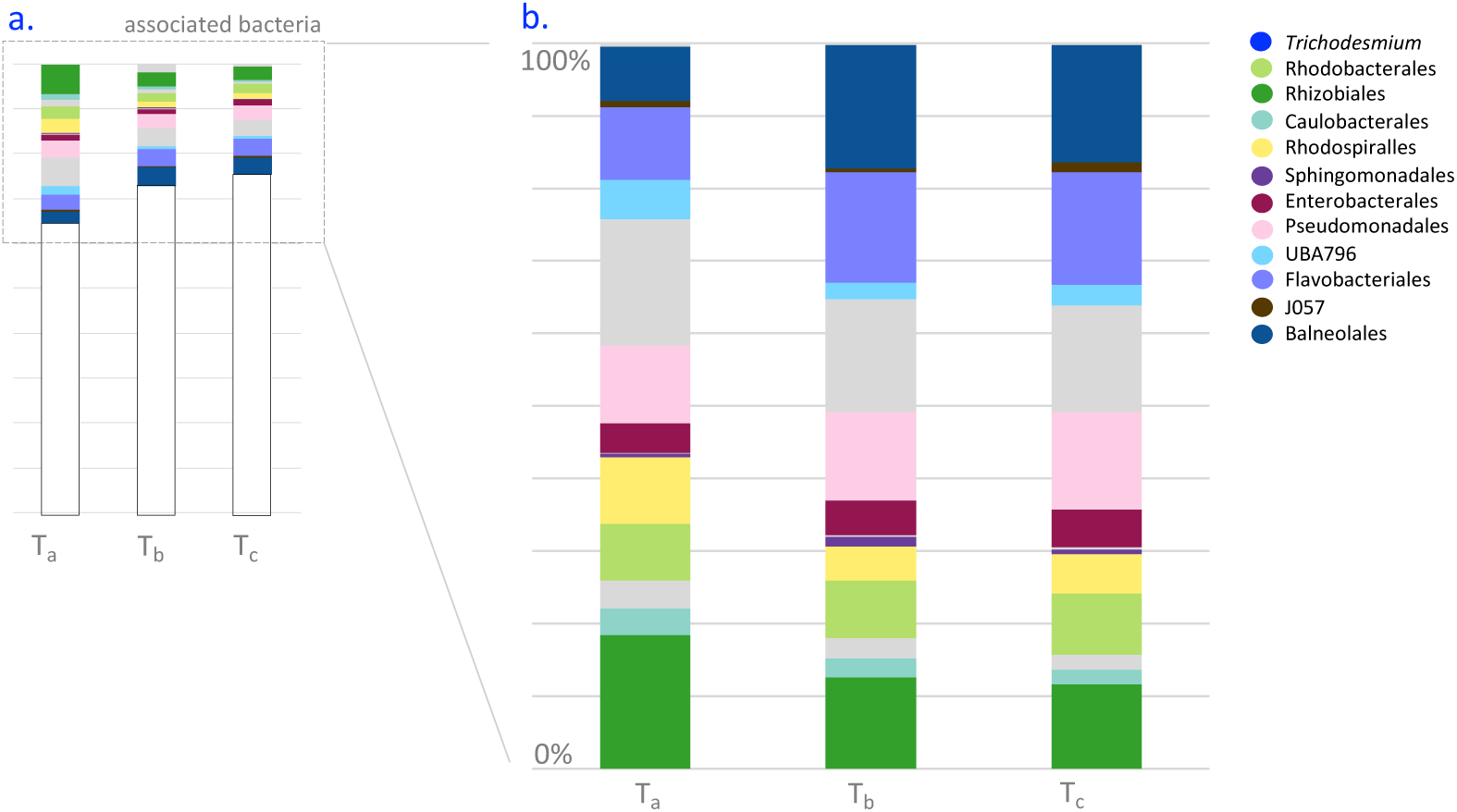
Relative abundance of bacterial MAGs from sampled *Trichodesmium* colonies indicate the associated bacterial distribution is consistent. The 51 MAGs are stacked, and colors reflect the different bacterial orders each MAG belongs to and ‘other’ in grey. The exact relative abundance value (%) for each MAG and its standard deviation can be viewed in Supplementary Table 1.

**Supplementary Figure 2.**
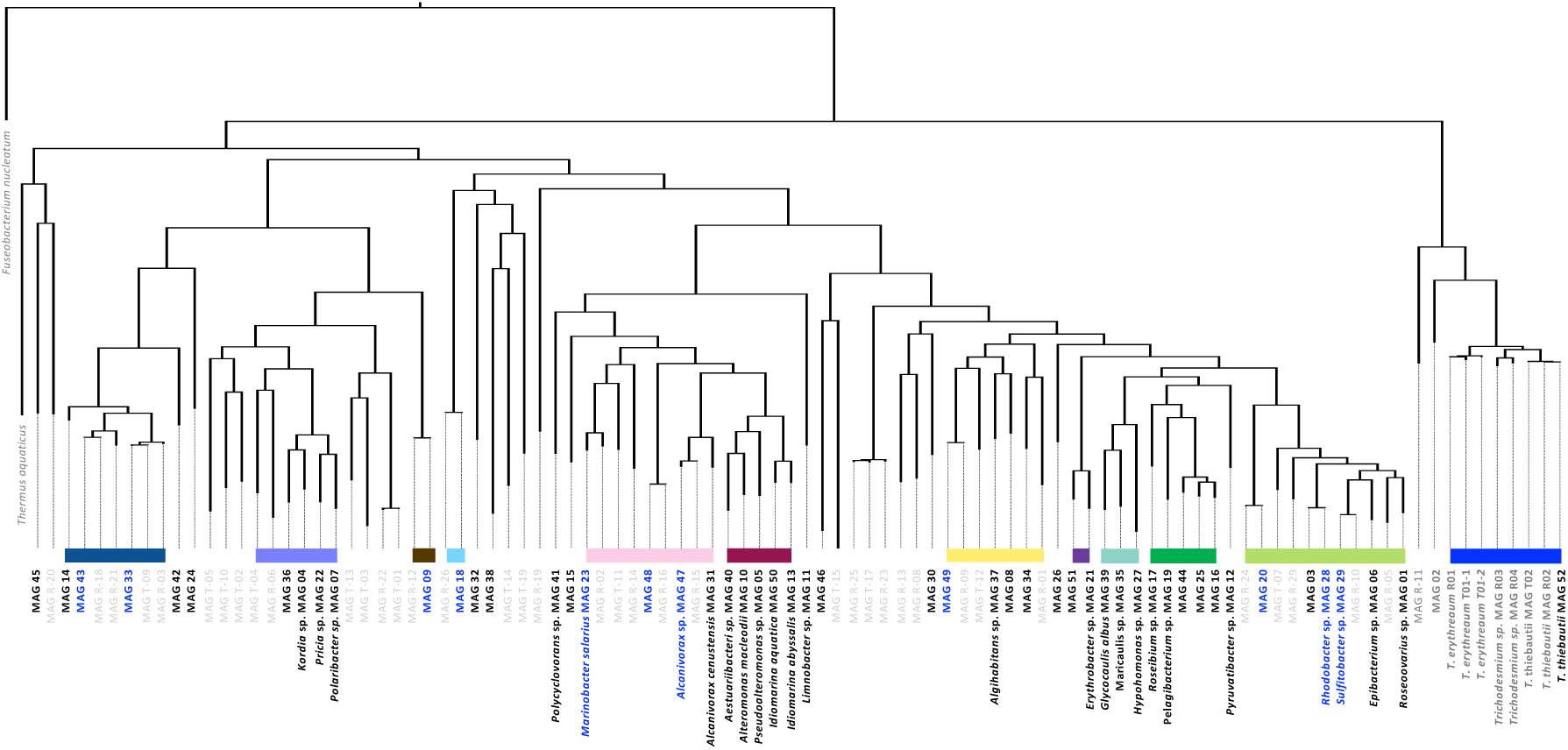
Phylogenetic tree of the 52 MAGs together with MAGs assembled from 3 other Trichodesmium metagenomic datasets from colonies collected in the Red Sea (28), the Pacific (12, 17) and the Atlantic (13) Oceans. MAGs from the 3 other datasets are highlighted in grey. MAGs deriving from samples of the Atlantic or Pacific Ocean are listed as MAG T-XX and those from the Red Sea are listed as MAG R-XX. The 11 MAGs that matched to those assembled from previous metagenomic datasets, using a 97.5 % identity cutoff, are highlighted in blue. Known bacterial orders are marked using different colored squares. The tree is rooted by the outgroup *Fuseobacterium nucleatum*. The full taxonomic description of each MAG in this dataset and others can be found in Supplementary Table 4.

**Supplementary Figure 3.**
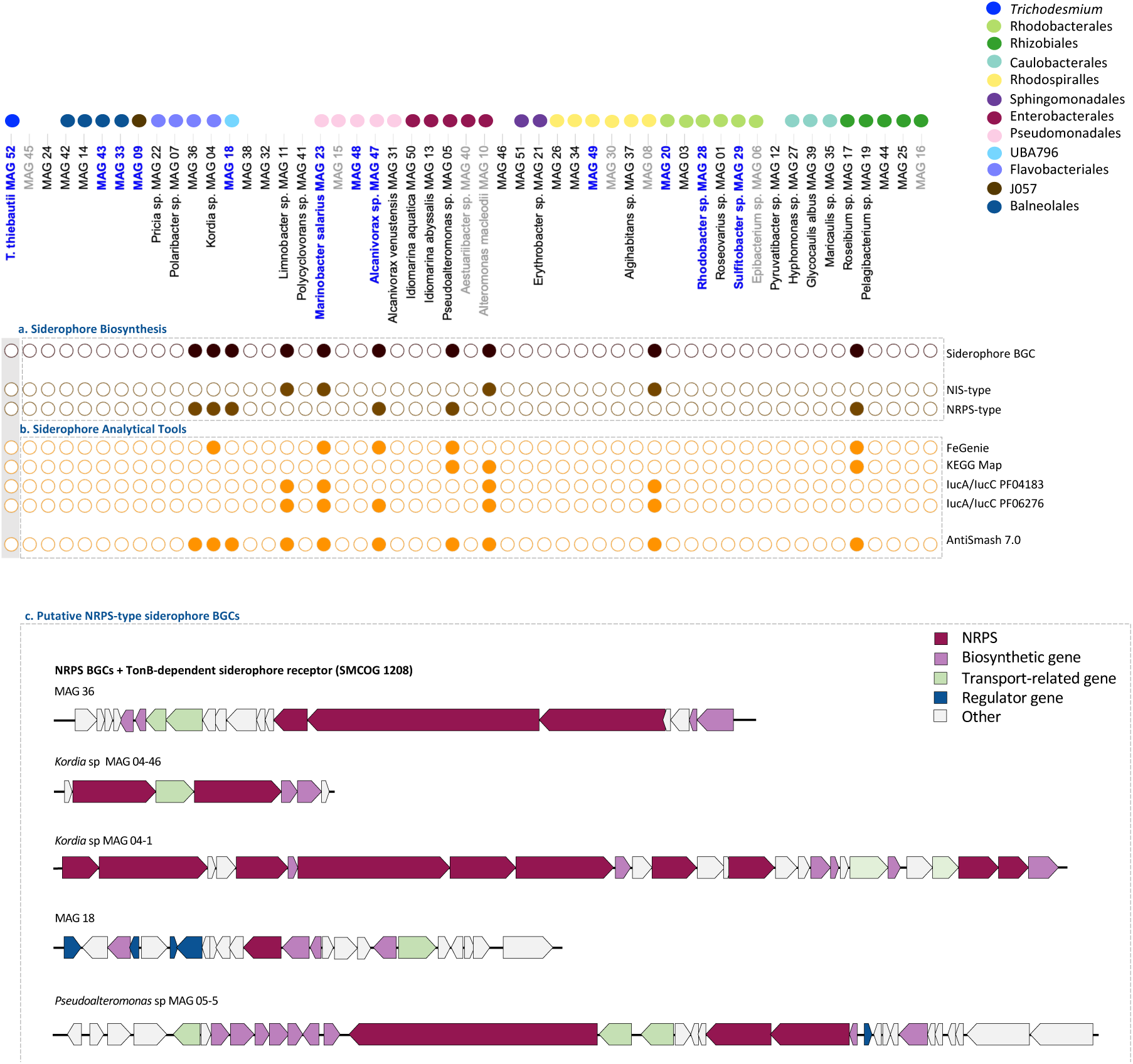
(a) NIS and NRPS-type siderophore biosynthesis pathways identified in MAGs of associated bacteria. (b) The siderophore BGC analysis tools used to probe and identify the different siderophore BGCs. Most tools are limited in their ability to discover siderophore BGCs while antiSMASH 7.0 was the most conclusive but also time-consuming (53). **(c)** Putative NRPS-type siderophore BGCs that did not match to a known siderophore within the MIBiG database but did containing a TonB-siderophore receptor (SMCOG 1208). This made us speculate that these are putative siderophores whose identity remains to be resolved.

**Supplementary Figure 4.**
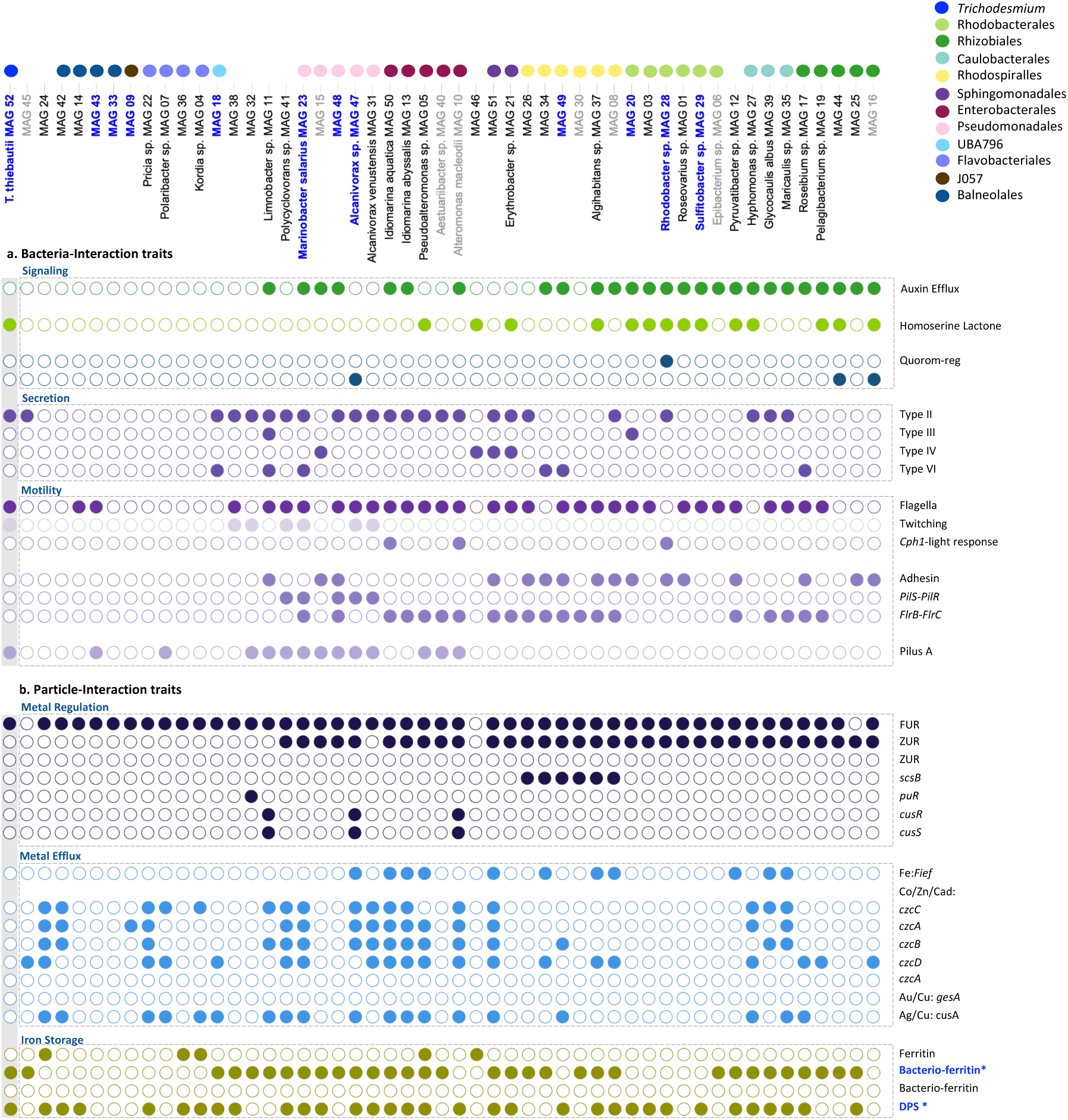
(a) MAGs were probed for bacterial-interactive traits. (68). MAGs from the phyla Proteobacter, contain the presence several interactive traits, including quorum sensing, gliding motility, adhesion, cell-to-cell signaling, and antibiotic resistance marker genes. Intriguingly, both the Trichodesmium MAG and several Proteobacter MAGs (e.g. Rhodobacter) contain genes for homoserine lactones (HLs), which are putative quorum sensing molecules. **(b) MAGs were probed for particle-interactive traits** (69). In comparison to *T. thiebautii* MAG 52, many MAGs from associated bacteria contained a wide variety of heavy metal efflux systems and metal-transcription factors. While *Trichodesmium*, like most Cyanobacteria, is known to be sensitive to heavy metals, our results indicate that the consortium is attuned towards an environment rich in metals, including particulate dust. Genes for each trait are noted in Supplementary Table 5

